# Biodiversity modulates the cross-community scaling relationship in changing environments

**DOI:** 10.1101/2025.03.07.642014

**Authors:** Vojsava Gjoni, Florian Altermatt, Aurélie Garnier, Gian Marco Palamara, Mathew Seymour, Mikael Pontarp, Frank Pennekamp

## Abstract

Organismal abundance tends to decline with increasing body size. Metabolic theory links this size structure with energy use and productivity, postulating a size–abundance slope of –0.75 that is invariant across environments. We tested the robustness of this relationship across gradients of protist species richness (1–6 species), temperature (15–25 °C), and time. Using replicated microcosms, we provide an empirical test of how temperature and biodiversity jointly shape the cross-community scaling relationship (CCSR). While our results support the expected of slope –0.75, we also found interactive effects showing the relationship is not invariant. Warming altered abundance scaling with size depending on richness; in high-richness communities, temperature favoured small protists, steepening the CCSR slope. These context-dependent responses emerged over time, suggesting a role of size-dependent species interactions in shaping responses to environmental change. Our findings demonstrate that cross-community size scaling is not fixed but shifts dynamically with ecological context.

## INTRODUCTION

Across ecological communities, smaller species tend to occur in higher numbers than larger ones. This consistent inverse relationship between body size and abundance is a well-documented empirical pattern (Damuth 1991; White *et al*. 2007), with a typical slope of approximately –0.75 on a log-log scale (Brown *et al*. 2004; Nee *et al*. 1991). The widespread occurrence of this pattern has made it central to community ecology, offering insights into how body size structures populations and communities (Arim *et al*. 2011; Enquist *et al*. 1998; Giometto *et al*. 2013; Jacquet *et al*. 2020; Li 2002; Long & Morin 2005; Meehan *et al*. 2004; Morán *et al*. 2010; White *et al*. 2004, 2007).

Beyond its descriptive power, the size–abundance relationship can link individual-level traits to ecosystem-level processes, serving as a tool for predicting how communities respond to environmental change (Gjoni *et al*. 2023; Gjoni & Glazier 2020). A central theoretical explanation for this pattern is the “energy equivalence”, which posits that total population energy use remains constant across body sizes (Brown *et al*. 2004; Damuth 1981). This is based on the observation that population abundance declines with body mass (∝ mass^-0.75^), while individual metabolic rate increases with mass (∝ mass^0.75^), resulting in a size-invariant energy flux per unit area. This assumption is grounded in the idea that population abundance is limited by resource availability, resulting in a trade-off: species with greater per capita energy demands must occur at lower densities. An increase in resource input in-turn could lead to a proportional increase in population abundance or sustain the same number of individuals but with increased energy demand (Isaac *et al*. 2011).

While theoretical models predict consistent size-abundance relationships, mounting empirical evidence suggests empirical patterns may be more variable and context-dependent. Body size and abundance are responsive to biotic and abiotic factors, which may lead to changes in the size-abundance scaling when either the abundance of small or large organisms is particularly affected (Gjoni *et al*. 2023). Indeed, deviations from the expected -0.75 slope for the size-abundance have been observed (Arim *et al*. 2011; Gjoni *et al*. 2017, 2023; Gjoni & Basset 2018; Gjoni & Glazier 2020; Long & Morin 2005; Meehan *et al*. 2004), suggesting that energy used may not be the same across all organism sizes (Nee *et al*. 1991) and size-abundance scaling is not as invariant as assumed.

Temperature represents one of the most pervasive environmental factors that could disrupt these scaling relationships, yet its effects remain poorly understood. Warming is predicted to accelerate metabolic rates, increasing the energy required to sustain individuals following the metabolic theory of ecology (MTE) (Brown *et al*. 2004; Gillooly *et al*. 2001). As a result, under constant energy availability, warming reduces carrying capacity by limiting the number of individuals that can be supported (Allen *et al*. 2002). However, because MTE assumes that temperature affects organisms similarly regardless of body size, it predicts that the slope of the size–abundance relationship should remain unchanged with warming (Fig. 1a). Contrary to this prediction, empirical studies often report deviations from the expected –0.75 slope in size–abundance relationships under warmer conditions (Gjoni *et al*. 2023; Morán *et al*. 2010; Pomeranz *et al*. 2022; Saito *et al*. 2021). One potential contributing factor is the temperature–size rule, which posits that organisms tend to attain smaller body sizes at higher temperatures (Atkinson 1995; Daufresne *et al*. 2009; Finkel *et al*. 2010; Gardner *et al*. 2011; Sheridan & Bickford 2011). However, the temperature–size rule alone does not necessarily explain changes in the slope of size–abundance relationships. If all organisms shrink proportionally, both small and large taxa could increase in abundance equally, maintaining the original scaling. A change in the slope would require a differential effect of temperature across size classes—where, for instance, smaller organisms benefit more from warming than larger ones, or vice versa. In addition, ecological processes not accounted for by metabolic theory, such as species interactions, may also contribute to observed deviations. These interactions can be sensitive to both temperature (Burnside *et al*. 2014) and body size (Freckleton & Watkinson 2001), further complicating predictions based solely on metabolic scaling principles.

**Figure 1.**
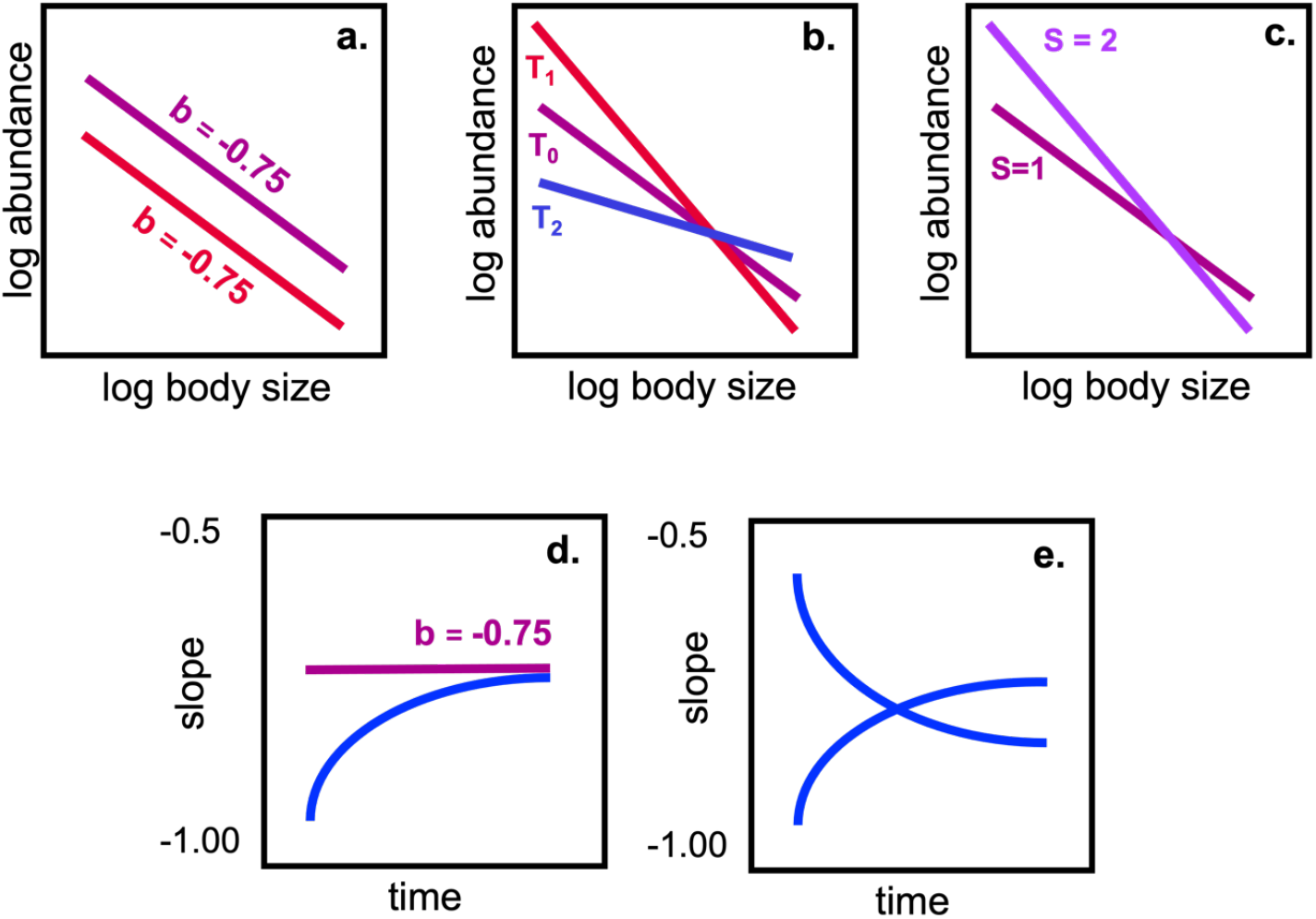
Conceptual overview of cross-community scaling relationships across temperature and diversity. a) Temperature and diversity may affect abundance and size regardless of the size of individuals, which may lead to changes in the intercept, but not the scaling. In contrast, smaller or larger individuals may be favoured at b) higher (T1) or lower temperature (T2), or c) lower or higher species richness, leading to changes in the cross-community scaling. Over time, the cross-community scaling relationship may be stable or converge to the expected slope of -0.75 (d) or show more variable trajectories with endpoints that differ from the expected slope (e).

Biodiversity can play a critical role in shaping the size-abundance relationship due to species interactions that determine community resource use and structure. While predation is a major focus of studies investigating the size structure of communities, competitive communities have been less frequently studied in the context of size-abundance scaling *(but see Long et al. 2006; Long & Morin 2005)*. Research on biodiversity-ecosystem functioning has, for example, revealed that increased species richness consistently leads to higher community abundance (Hooper *et al*. 2005; Tilman *et al*. 2001). Higher community abundance can be driven by highly productive species being more likely to occur in species-rich communities (i.e., selection effects), or resource partitioning (i.e., complementarity) among species, reducing competition (Blackburn & Gaston 1996; Tilman *et al*. 2001). How community size structure is related to biodiversity effects is less well understood. Size variation in competitive communities is often small (Tilman *et al*. 2004) and size structure is rarely linked to complementarity or selection effects (but see Norkko *et al*. 2013). This suggests size-abundance scaling is unaffected by species richness per se (Figure 1a).

Temperature and diversity can also interactively shape the size-abundance relationship, if they disproportionally affect the interactions between larger and smaller organisms (Fig. 1b). Higher temperatures tend to reduce the number of larger individuals, due to their higher metabolic demands, creating a competitive advantage for smaller individuals due to their lower resource requirements (Petchey et al. 1999). Temperature may also counteract some of the positive biodiversity effects observed at higher richness due to direct negative physiological effects on biomass, leading to shallower or even negative biodiversity-abundance relationships (Parain *et al*. 2019; Pennekamp *et al*. 2018). In addition to these metabolic constraints, increased species richness can lead to reductions in population-level density due to interference competition, niche overlap, or antagonistic interactions among co-occurring species (DeLong 2021; Finke & Denno 2005; Turchin 2003). These negative diversity– density relationships may also alter community size structure, especially if the strength or symmetry of competition varies with body size. Larger organisms are often competitively superior, as increased richness intensifies asymmetric resource competition (Lawton & Hassell 1981; Gómez & González-Megías 2002). Yet, whether larger or smaller species are favoured depends on the ecological and thermal context. Notably, in the absence of an interaction between activation energy and body size, the metabolic effects of temperature scale proportionally across sizes—meaning temperature accelerates the metabolism of many small organisms and a few large organisms to the same relative degree, with the proportional effect governed by –E/kT (Brown *et al*. 2004). Therefore, temperature alone does not necessarily alter the slope of density–mass scaling unless body-size-dependent differences in thermal sensitivity are present. In species-rich communities, such differential sensitivities—amplified by complex interactions—may shift the competitive balance, causing richness to drive either steeper or shallower size-abundance slopes depending on which size classes gain a relative advantage (Fig. 1c). More generally, temperature and biodiversity can affect body size distributions and abundance through a suite of interconnected mechanisms. These include: shifts in metabolic rates and energy demand with temperature; differential thermal sensitivities among species, which alter interaction strength and performance; resource partitioning or competition intensity in diverse communities; and the tendency of larger organisms to dominate in systems with strong interference effects. By altering these underlying processes, temperature and diversity may interactively drive changes in both the intercept and slope of the size–abundance relationship, particularly when interactions between these factors amplify size-structured outcomes.

Effects of biotic and abiotic factors on the size-abundance scaling may further vary through time, since ecological communities are complex, adaptive systems which exhibit temporal dynamics in response to both biotic and abiotic factors (McGrady-Steed *et al*. 1997). Over time, changes in temperature and biodiversity may alter the relative proportions of organisms of different sizes within a community. Due to the equilibrium assumptions of the energy equivalence (Brown *et al*. 2004; Nee *et al*. 1991), we would expect the slope to converge to -0.75 in the long run. Indeed, experimental studies have shown that closed communities with controlled species richness (e.g., low vs. high richness) converge to a stable size-abundance relationship (Fig. 1d; Long & Morin 2005). In contrast, natural communities can exhibit fluctuating size-abundance relationships over time, driven by dynamic species interactions and environmental variability (Fig. 1e; White *et al*. 2004) that deviate from the expected slope.

Here, we investigated the size–abundance relationship across temperature and biodiversity (i.e., initial species richness) in controlled microcosms over time (Pennekamp et al. 2017, 2018). All communities were initially started from the same set of species, thus changes in size and abundance were integrating both phenotypic change in size and abundance. Given evidence that temperature can differentially affect organisms of different sizes, and that species richness may mediate these effects through competitive interactions, our hypotheses explicitly test whether these factors alter size– abundance scaling. Based on a null model informed by the MTE, we hypothesize the following: (1) the size–abundance relationship initially follows a slope of –0.75, as predicted by MTE, (2) this slope may vary with temperature and species richness due to differential effects on small and large species, rather than remaining invariant, and (3) the slope may converge toward –0.75 over time as communities stabilize.

While ecological theory has traditionally emphasized within-community size spectra, which describes the distribution of individual body sizes and abundances within a single community at a given time, our study takes a broader perspective. We examine cross-community scaling relationships (CCSRs), which link the average body size of individuals to the total community abundance across different communities (White *et al*. 2007). While size spectra offer a useful summary of internal community structure, they do not retain the direct relationship between size and abundance necessary to assess ecological scaling patterns. In contrast, CCSRs explicitly link body size to total abundance at the community level and are therefore well suited for testing predictions based on energy equivalence and metabolic scaling theory (Isaac *et al*. 2011). Our experimental design included 53 protist communities per temperature treatment, each varying in initial species richness and composition. By comparing CCSRs across temperature and richness gradients, we were able to examine how warming alters the scaling of abundance with size—particularly whether these relationships deviate from the theoretical –0.75 slope expected under the metabolic theory of ecology. To further understand the role of species interactions on the CCSR, we created polycultures without interactions by pooling monoculture data with the same richness and composition (i.e., communities with no interaction expectation) as the real polycultures that we studied experimentally. Unlike previous observational studies, this controlled approach allowed us to directly test how temperature and diversity jointly shape cross-community scaling.

## MATERIALS & METHODS

We used microcosms to test the effect of temperature and species richness on the CCSR. To this end, we factorially manipulated temperature (15, 17, 19, 21, 23, and 25 °C) and species richness (1 to 6 species) in communities of six bacterivorous ciliate species differing in average cell volume (ordered from the smallest to the largest): *Dexiostoma campylum (0*.*0000109 millicubicmeter), Tetrahymena thermophila (0*.*0000173 millicubicmeter), Loxocephalus* sp. *(0*.*0000542 millicubicmeter), Colpidium striatum (0*.*000111 millicubicmeter), Paramecium caudatum (0*.*000180 millicubicmeter), and Spirostomum teres (0*.*000488 millicubicmeter)*. The species were cultured in organic protist pellet medium on which the freshwater bacterium *Serratia fonticola* was inoculated as the sole protist food source. Due to the non-sterile handling of cultures, additional bacteria may have colonized the microcosms throughout the experiment. The organic medium consisted of protist pellets (Carolina Biological Supplies, Burlington, NC) at a concentration of 0.55 g per L of Chalkley’s medium, and two wheat seeds for slow nutrient release (for details on cultivation and experimental procedures, see also Altermatt *et al*. (2015)). Stocks of protist cultures (monocultures) were established before the start of the experiment to inoculate the microcosms. Single species cultures (richness = 1) were started at a density of three individuals per mL in 100 mL medium kept in 250 mL Duran bottles. Multispecies communities with species richness ranging from 2 to 6 species were initiated as a fixed fraction of the species-specific carrying capacity. For example, 2-species communities were started with 20% of their respective carrying capacity, while 4-species communities were started with 10% each. While having the same fraction of carrying capacity, smaller and more abundant species started with a higher number of individuals than larger and less abundant species. Since each species occurred the same number of times in each richness level, there was no bias in terms of the number of large or small species across richness levels. Since communities were closed to dispersal, richness can only decrease, and species may be lost due to extinctions. We replicated each level of species richness and composition at least twice for all levels including an additional replicate for the lowest and additional three replicates for the highest levels of species richness resulting in 115 experimental units per temperature. Communities were randomly assigned to their temperature level and placed in climate-controlled incubators (Pol-Eko Aparatura, Wodzislaw, Poland) set to 15, 17, 19, 21, 23, or 25 °C. We used two independent incubators for each temperature level. For an overview of the experimental design, refer to table S1.

To measure abundance and size of individual cells, we sampled each experimental unit every day for the first 7 days, then 3 times per week for the following 50 days, and a final sampling 7 days later. We used an established automated video species classification analysis (Pennekamp *et al*. 2015, 2017) to calculate the average size (i.e., volume) and abundance of each species. Since monocultures and polycultures had different starting densities, we aligned the time series to make the dynamics directly comparable. Please refer to Supplementary material section S1 for details. We then calculated the cross-community scaling relationship relating total community abundance (sum of the abundance of all species in the community) and the weighted mean size of an individual in the community (White *et al*. 2007).

We used a regression approach to quantify the relationship between abundance (response) and size, richness and temperature in our controlled microcosm communities. We scaled size, temperature and richness treatments (by subtracting the respective mean from all values) to facilitate the interpretation of model coefficients (Schielzeth 2010). The random structure for all models consisted of a random intercept for community composition and a random intercept for incubator. We then perform four analyses: 1) we estimated the overall cross-community scaling relationship across protist communities, regressing total abundance on mean individual size (White *et al*. 2007). 2) We regressed total abundance on mean individual size, temperature, and richness as well as all interactions between main effects averaged over time. We included all possible interactions of the 3-way interactions models (cell volume x temperature x richness). 3) We fitted the previous model (3 main effects and all possible interactions, same random effects) to each time point to understand how estimates change through time.

4) To understand the role of species interactions on the CCSR, we generated communities with no interaction expectation by pooling monoculture data with the same richness and composition as the real polycultures that we studied experimentally. We then used the same model as 3) to compare the effects of size, richness, and temperature, as well as their interactions between the real communities and communities with no interaction expectation. Deviations in coefficient estimates would be putatively driven by species’ interactions. All analyses were performed in the statistical computing environment R (version 3.6.1).

## RESULTS

### Interacting temperature and richness effects on CCSR

Averaging over time, we found a strong negative CCSR (*b* = -0.71, CI = -0.777 to -0.644) across community richness and temperature treatments, undistinguishable from the expected value of *b* = - 0.75 (Fig. 2A). Although deviations from the linear fit are visible, they disappear when data was stratified by richness (Fig. 2B).

**Figure 2.**
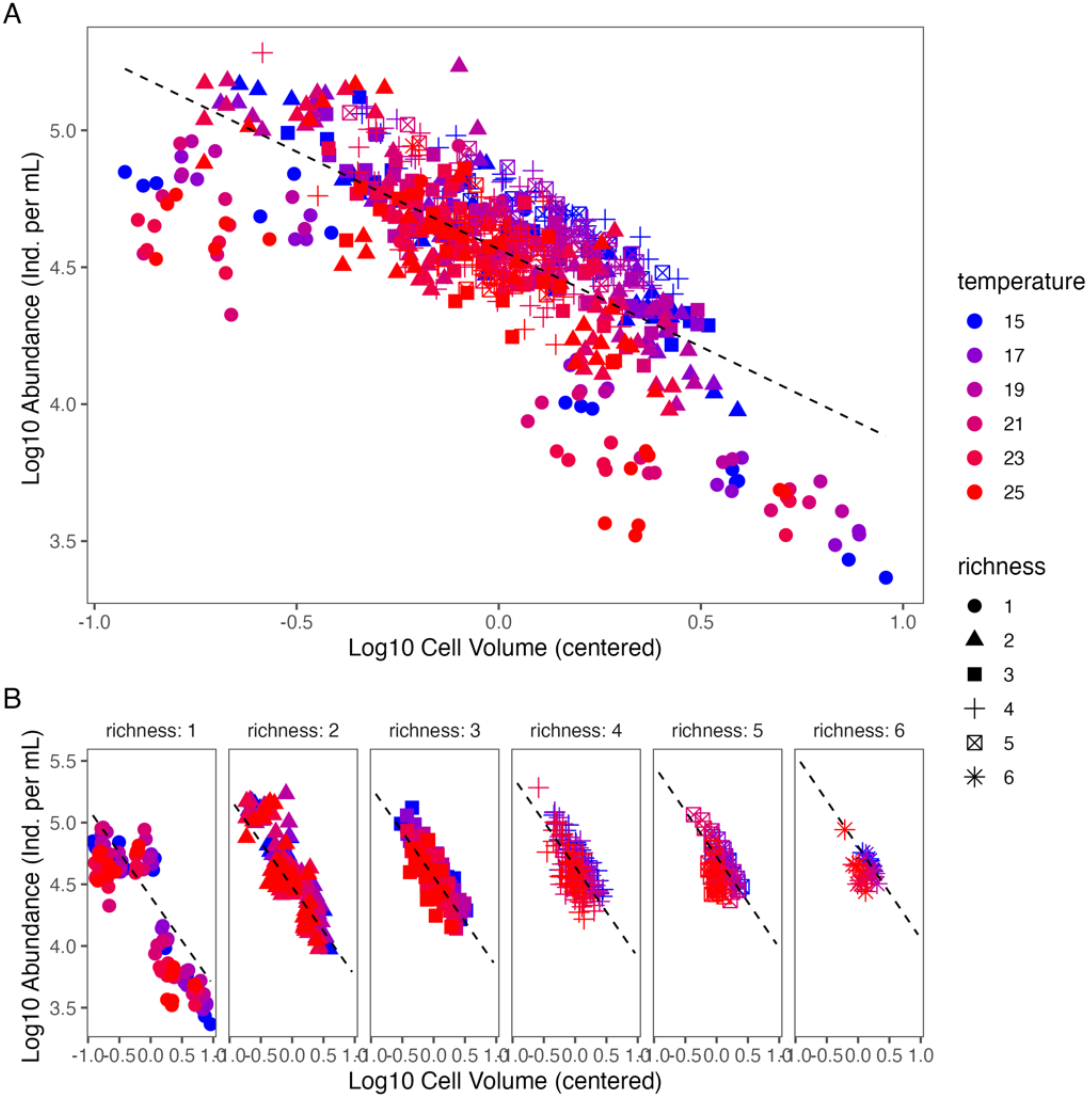
The relationship between centred average log10 cell volume (μm μL^-1^) and average log10 total abundance of protist communities, approximated with a simple linear fit (A). Deviations are visible at the lowest (< -0.5 log cell volume) and largest average cell sizes (> 0.5 log cell volume) but disappear when stratified by richness (B).

Including temperature and richness in the model (Table 1), the CCSR slope (*b* = -0.749 CI = - 0.813 to -0.685) perfectly matched the expected value. However, temperature and richness jointly changed the effect of size on abundance (volume:temperature coefficient = -0.015, CI = -0.024 to - 0.005 and volume:richness coefficient = -0.055, CI = -0.100 to -0.010). The effect of size on abundance was furthermore mediated by a three-way interaction (volume:temperature:richness coefficient = - 0.012, CI = -0.018 to -0.007). In other words, at low richness, the abundance of small and large protists was not affected by temperature (similar slope), whereas in more species rich communities, temperature increased the abundance of smaller protists relative to larger ones, resulting in a steeper CCSR (Fig. 3, Figure S1, Table 1). Importantly, these changes were not driven by extinctions of the larger species (see sections S4). Using only two replicates for each richness level (leading to a balanced design) yielded quantitatively and qualitatively similar results.

**Table 1.**
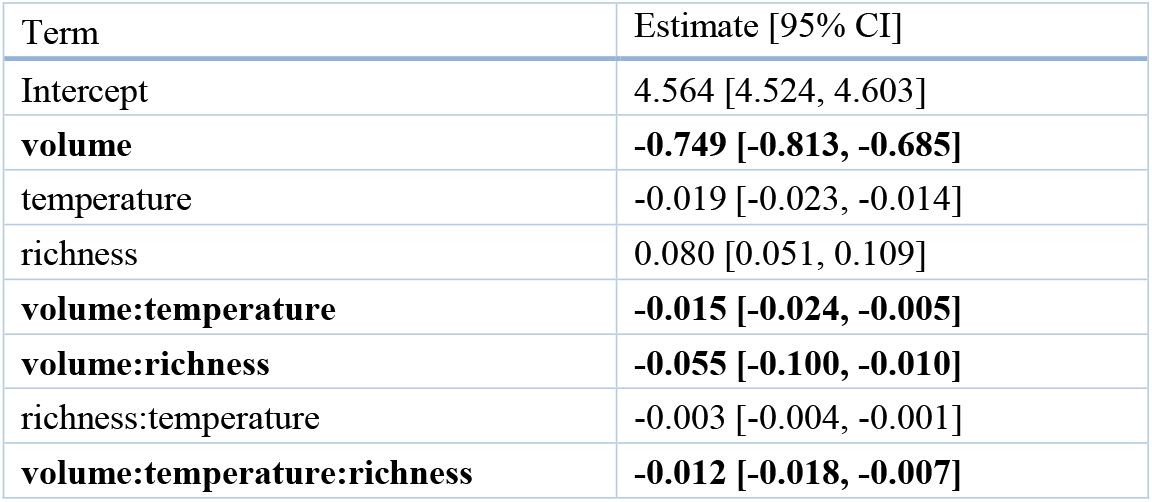
Temperature and species richness had an interactive effect on CCSR across protist communities. A confidence interval (CI) not overlapping with zero indicates a significant effect. All significant terms including the effect of volume on abundance are shown in bold. All predictors were mean centred for the analysis.

**Figure 3.**
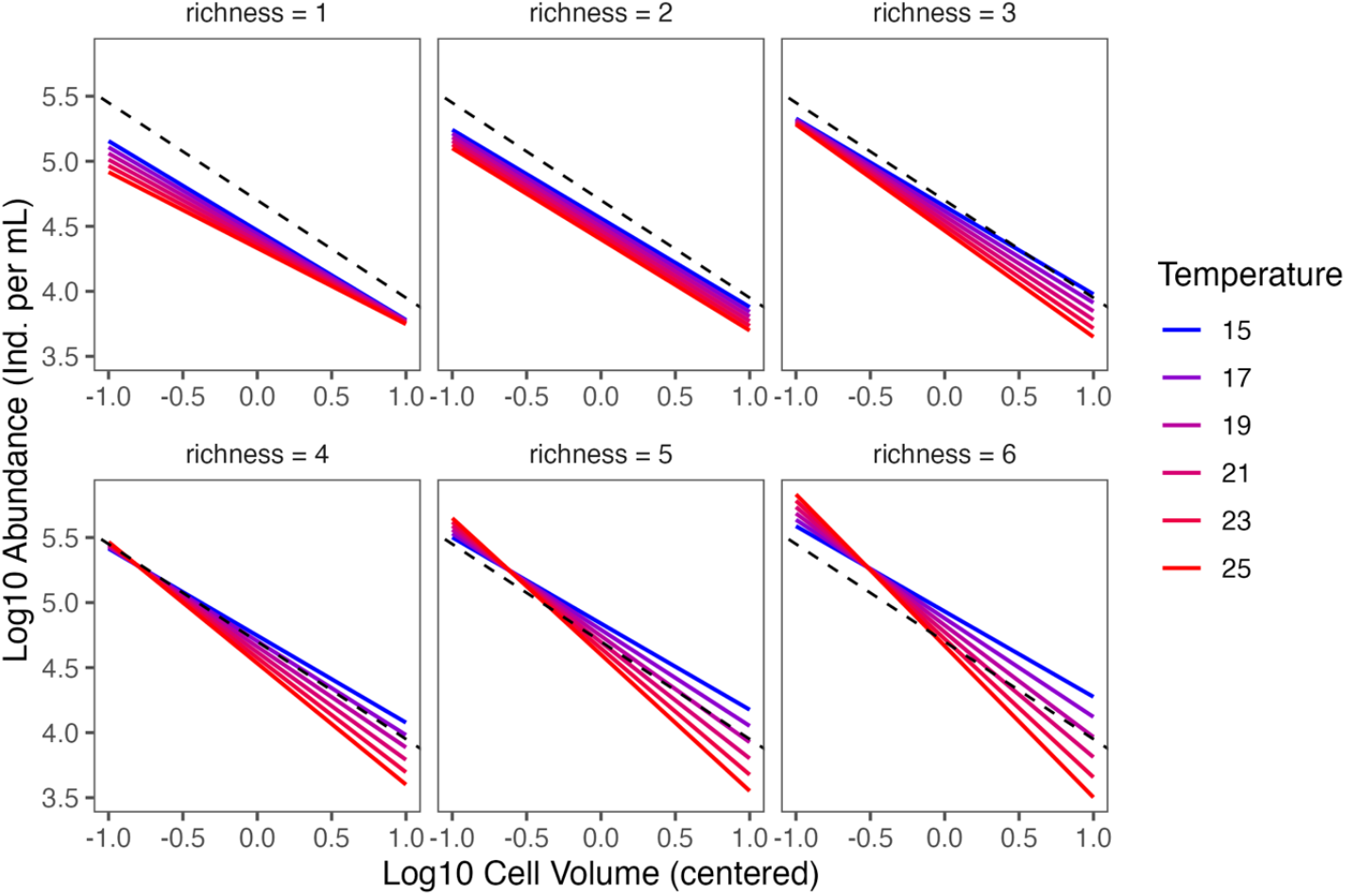
Temperature and richness interact to alter protist community CCSR. The dashed line indicates the expected size-abundance scaling of -0.75. The figure shows the fitted size-abundance scaling based on the model presented in Table 1. A slope steeper (shallower) than -0.75 indicates that total abundance declines faster (more slowly) than expected with increasing average cell volume of the community.

### CCSR through time

At the beginning of the experiment (Fig. 4A), CCSR was steeper (*b* = -0.9) than expected but stabilized on day 20 with a slope of b=0.75, which was expected. Afterwards the CCSR slope became shallower than expected (*b* = -0.35). Similar temporal patterns were observed when we used community evenness accounting for species extinctions and changes in relative abundance (see Section S3). Initially, communities with smaller average individual sizes tended to have higher abundance compared to those with larger individuals. Over time, communities with intermediate species richness (e.g., 5–7 species) showed relatively stable or slightly increasing abundance trends. By the end of the experiment, communities with larger average individual sizes had higher abundance than would be expected based on early patterns.

**Figure 4:**
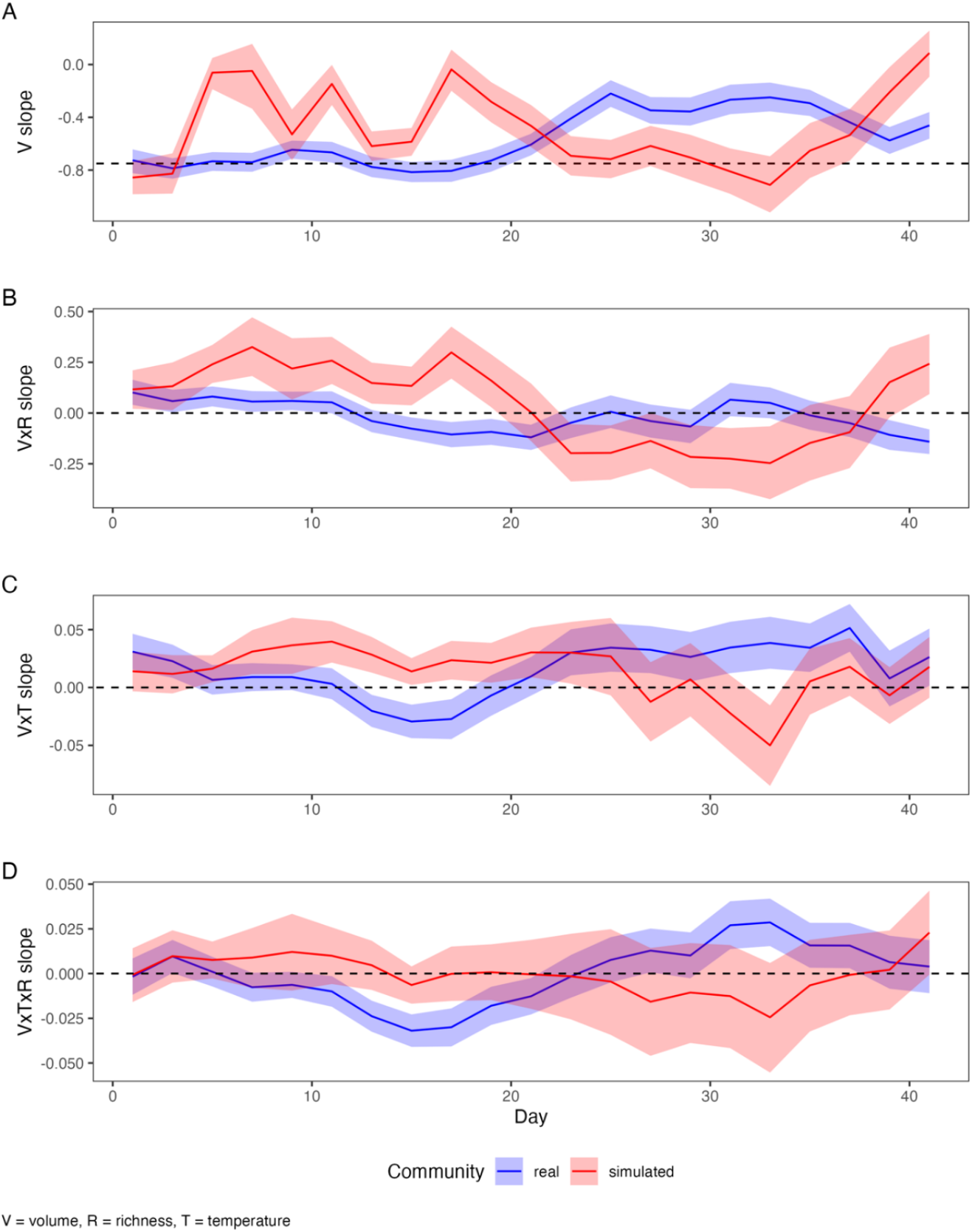
Temporal dynamics of effect sizes (and 95% confidence intervals) of a linear mixed-effects model where abundance is modelled as a function of size (V = volume), temperature (= T) and richness (= R) and their interactions for the observed and communities with no interaction expectation (in blue and red, respectively). The x-axis shows the day. A) Effect of cell size (V) on abundance, B–C) effect of two-way interactions between size and temperature (V x T), size and richness (V x R) on abundance. D) Three-way interaction between size, temperature and richness (V x T x R) on abundance. The dashed lines indicate a slope of -0.75 in panel A and zero in panels B–D (a deviation from zero would indicate a slope modification). A slope steeper (shallower) than -0.75 indicates that total abundance declines faster (more slowly) than expected with increasing average cell volume of the community.

Temporal analysis of the CCSR revealed dynamic patterns: temperature (V x T) initially favored larger organisms but shifted to favor smaller ones after day 17, becoming non-significant by the experiment’s end. Species richness (V x R) initially reduced cell size effects but reversed after day 20 to promote larger organisms. The temperature-richness interaction (V x T x R) alternated between negative and positive effects throughout the experiment, resulting in high richness communities being more strongly affected by temperature than low richness communities.

Comparing real polycultures with communities having no interaction expectation revealed contrasting temporal dynamics (Fig. 4). Initially, no-interaction communities showed steeper size-abundance slopes than expected, indicating higher relative abundance of smaller organisms. After day 14, these slopes became shallower (approaching -0.40 by day 20) while real communities maintained - 0.75 until day 20. By day 30, no-interaction communities returned to expected values while real communities became increasingly shallow. Unlike observed protist communities, the combined temperature-richness effect showed no significant influence, indicating that temperature’s impact on size-abundance slopes was not richness-dependent in the no-interaction scenario.

## DISCUSSION

Our study revealed that the cross-community scaling relationship of protist communities is mediated by temperature, biodiversity (i.e., initial species richness), and time, challenging the widely held assumption that CCSRs remain stable within a trophic level across time and space (Damuth 1981; Enquist & Niklas 2001; Niklas & Enquist 2001). We found a negative CCSR (b ∼ -0.75) across richness, and temperature variation, in line with the expected slope of the energy equivalence (Damuth 1991; Nee *et al*. 1991), confirming previous studies using the CCSR across phytoplankton (Li 2002), macroinvertebrates, amphibians, and fish (Arim *et al*. 2011), rodent (White *et al*. 2004) and tree communities (Enquist *et al*. 1998).

A CCSR is close to -0.75 is sometimes mirrored by a metabolic rate with a 0.75 scaling exponent with mass (Brown *et al*. 2004), for instance in phytoplankton communities (Malerba *et al*. 2017; Perkins *et al*. 2019). Despite this support of the –0.75 slope, deviations are frequently reported in within-community or population-level studies, where scaling relationships may vary depending on the level of organization, sampling method, or ecological context (DeLong *et al*. 2010; Hatton *et al*. 2019; White *et al*. 2007). An experimental study using protist communities initially divided into low and high species richness showed no change over time with a cross-community size scaling of -1, which converged over time (Long & Morin 2005). As we lack direct measurements of metabolic rates for our focal protist species, we cannot confirm whether the classic metabolic scaling holds in this system.

Our experiment revealed a CCSR close to -0.75, suggesting that the scaling of metabolic rate with body size may be a possible explanation. However, when we explicitly considered the abiotic and biotic context of communities, we found that the CCSR of protist communities is mediated by temperature, biodiversity and their interaction. Interacting effects between CCSR and temperature as well as diversity, are inconsistent with energetic equivalence. In our high-richness communities, small organisms were favored (more abundant than expected) with high temperature, whereas large organisms were favored (more abundant than expected) when the temperature was low. Smaller organisms have lower absolute resource requirements (Gillooly *et al*. 2001; Reuman *et al*. 2014), allowing for a higher number of smaller organisms to be sustained (Li *et al*. 2006; Morán *et al*. 2010). Temperature may lead to smaller organisms on average according to the temperature-size rule (Atkinson *et al*. 2003), however, this physiological effect will affect organisms regardless of their size. Therefore, this cannot explain the observed changes in CCSR, requiring additional explanations. Species richness increased abundance, which is in line with experimental evidence that increasing the number of species (i.e., in this case, competing for the same resources, such as bacteria) increases the total acquisition of resources due to complementarity (Gómez & González-Megías 2002b). However, richness did not interact with size, suggesting that complementarity and selection effects are not size-dependent in our communities. We propose that temperature-dependent changes in species interactions provide the key explanation.

The interactions of the CCSR with temperature and richness may hence be driven by temperature-dependent size asymmetries in resource competition (Cotgreave 1993; Nee *et al*. 1991; Russo *et al*. 2003). One possible explanation is that temperature alters the competitive asymmetry among organisms of different body sizes by differentially affecting their metabolic rates, feeding efficiencies, and growth responses—traits that govern resource acquisition and competitive ability. At higher temperatures, smaller organisms, with inherently faster metabolic rates, may outcompete larger ones by exploiting resources more quickly. Conversely, at lower temperatures, the metabolic and growth advantages of small size diminish, potentially allowing larger-bodied organisms to dominate through superior resource access or longer generation times. These patterns are consistent with size– temperature relationships described in the context of Bergmann’s rule (Blackburn et al. 1999).

Another plausible mechanism for the increasing effect of temperature with rising species richness is that temperature intensifies the strength and asymmetry of species interactions, particularly in more complex communities. In species-rich systems, where interaction networks are more densely connected, warming can amplify small differences in species’ thermal sensitivities, resulting in cascading effects on competition and resource partitioning (Dell et al. 2014; Gilbert 2014; O’Connor et al. 2011). These dynamics may shift dominance patterns and energy flow, thereby altering the CCSR more strongly under high richness. For instance, stronger interaction networks under warming can lead to greater resource monopolization by thermally favoured species, magnifying size asymmetries (Sentis et al. 2017; Uszko et al. 2017). Thus, species richness not only increases the potential for interaction-driven community structuring but also makes the system more sensitive to temperature-induced change.

Previous analyses on our protist communities have shown that species interactions become more asymmetric when temperatures rise. Tabi et al. (2020) showed that this leads to lower coexistence of protist communities and a decoupling between abundance in isolation and abundance in communities. Comparing the temporal dynamics of size-abundance scaling in communities with no interaction expectation and real communities provides indirect evidence for this: in observed polycultures, the initial slope aligned with the expected slope of -0.75, and over time, there was an interactive effect of size, temperature and richness. In contrast, this pattern did not occur in communities with no interaction expectation, as they lack the influence of species interactions. Maynard et al. (2020) estimated coefficients that describe temperature effects on species interactions, showing mostly negative effects of temperature and decreasing coefficients across the temperature gradient. Jointly, this suggests that the observed steepening of the size-abundance scaling in high richness communities at high temperature is driven by species interactions that are both temperature-dependent and size asymmetric.

Our results show that size-abundance scaling can shift dynamically over time. Contrary to expectations, the slope remained relatively stable in the first half of the experiment but deviated from the -0.75 benchmark by the end, indicating a greater proportion of larger individuals in the community. While this pattern diverges from the predictions of the Metabolic Theory of Ecology (MTE)—which suggests that energy use becomes size-independent over time (Brown et al. 2004)—it highlights that energy distribution in communities may become more complex, especially over extended timeframes than suggested by theory. Specifically, larger individuals may acquire more energy than smaller ones as communities mature, aligning with prior research suggesting resource monopolization by larger protists under resource scarcity (Malerba & Marshall 2019; Tabi et al. 2019). Their capacity to store resources may give them a survival advantage during periods of limited availability (Fenchel 1980). This idea is further supported by hypotheses proposing that larger organisms endure fasting conditions better due to their storage capacity (Millar & Hickling 1990), and experimental work showing that larger microbial cells maintain efficiency under shifting nutrient levels (Malerba et al. 2018; Marshall et al. 2022). Similar trends have been observed in field studies, where temporal changes in CCSRs were linked to shifts in species size and composition (White et al. 2004). However, given the arbitrary endpoint of the experimental timeline, we caution against overinterpreting the final size-abundance patterns as indicative of long-term outcomes.

Our study system differs from natural systems, which should be considered when interpreting the results. First, since we did not allow for colonization by warm-adapted species/genotypes, we cannot assess community turnover as a mechanism affecting community size structure under warming (Khaliq *et al*. 2024). Second, microcosms are closed systems that do not allow for emigration or metacommunity dynamics, which can buffer or balance ecological processes across space (Seymour *et al*. 2015). Although all microcosms initially received the same amount of abiotic resources and energy inputs were periodically replenished, the overall energy supply is not constant over time. Additionally, we did not quantify the bacterial communities and their dynamics, which may further influence ciliate community structure and interactions (Saleem *et al*. 2013). Microcosms can be colonized by air-borne bacteria; the chance of such colonization events increases through time and may have been affected by our time alignment. Replicate microcosms showed very repeatable dynamics, therefore we judge such chance events have negligible effects on our results. Therefore, temporal effects need to be judged cautiously and do not provide strong evidence against energetic equivalence. Third, we do not know whether the observed changes in cross-community scaling are due to phenotypic plasticity, or also include evolutionary responses. Despite these differences, our approach provides important complementary insights to analyses of cross-community size scaling in natural communities.

## CONCLUSION

Our experiments suggest that temperature and species richness play an important role in shaping the size structure of ecological communities through time. We hence caution against assuming that energy flux is size-independent in dynamic ecosystems, as changes in the biotic and abiotic environment may change community size structure and hence the flow of energy. Testing these assumptions with experimental communities provided new insights into the mechanisms driving community size responses to environmental change, improving our ability to predict how biodiversity may influence the size structure of communities in the face of global environmental change.

## ACKNOWLEDGEMENTS

VG was funded by a Swiss National Science Foundation Scientific exchanges fellowship (IZSEZ0_203498). We thank Roman Alther, Yves Choffat, Emanuel Fronhofer, Pravin Ganesanandamoorthy, Suzanne Greene, Katherine Horgan, Elvira Mächler, Thomas M. Massie and Owen Petchey for help with the data collection.

## Notes

### Competing Interest Statement

The authors have declared no competing interest.

### Summary of Updates

The manuscript has been revised to improve the clarity of the introduction and discussion, and provides additional analyses on the role of extinctions in driving patterns of cross-community scaling relationships.

