## Supplementary materials for "Biodiversity modulates the cross-community scaling relationship in changing environments"

### S1.1 Video microscopy and processing

The videos were taken using a stereomicroscope (Leica M205 C) with a 16-times magnification mounted with a digital CMOS camera (Hamamatsu Orca C11440, Hamamatsu Photonics, Japan). We used the BEMOVI package (Pennekamp *et al.* 2015, version 1.0.2) to process the 18,720 videos collected during the experiment and extract the raw trajectories. Global segmentation and tracking parameters were defined for the automated processing of videos (Pennekamp *et al.* 2017). The difference lag was defined to be 2 seconds, particle size was restricted to 5 to 2000 pixels, and the grey threshold was set to an intensity value of 10. For particle linking, we specified a link range of 3 frames and a displacement of 20 pixels. These settings were optimized using a subset of videos (spanning sampling dates of all single, two, and six species combinations at 15 °C, 21 °C, and 25 °C). Video settings were optimized to rather include false positives rather than exclude true positives at this step and exclude false positives later in the processing pipeline. For further details regarding video processing, please refer to (Pennekamp *et al.* 2017). After tracking, trajectories were filtered to remove artifacts such as spurious trajectories (e.g., floating debris). Trajectories for analysis were required to show a minimum net 7 displacement of at least 50 μm, a duration greater than 0.2 seconds, and a detection rate of 80 percent (for a trajectory with a duration of 10 frames the individual has to be detected on at least 8 frames), a median step length greater than 2 μm and a minimum mean speed of 50 pixels per second.

We applied machine learning (random forest classification) on quantitative properties such as cell size and shape to predict to which species an individual belongs. We thus identified individuals from all species automatically from the videos and quantified their cell size. Details of the procedure can be found in the supplement (Pennekamp *et al.* 2017) for a detailed description of the experimental design, setup of micro-ecosystems, and sampling regime.

### S1.2 Calculation of abundance and individual size

We calculated individual cell volume based on cell length and width measurements using the following equation: cell volume = 4/3 * pi * (width /2) ^ 2 * (length/2) (Laakso *et al.* 2003). Volume is reported in microlitre. Abundance was calculated by counting the number of individuals of a given species on the 50th frame of a video, thus avoiding double counting of individuals, and then upscaled to density per mL.

#### S1.3 Time series alignment

We only consider the data points of monoculture time series after reaching 20% of their carrying capacity, in correspondence with the polyculture time series as done in previous analyses (Pennekamp *et al.* 2018). The alignment did not influence the qualitative patterns observed over the full experimental period. Time series were cut to have a maximum length of 42 days. Alignment of mono- and polyculture time series led to differences in the time points where abundance and size were measured. We used cubic hermite splines to interpolate the mean abundance and mean volume of species in each experimental microcosm. We manually checked that interpolations closely agreed with the size and abundance dynamics observed. In rare cases, when either negative abundances or volumes resulted from the interpolation, we discarded the observation. For all subsequent analyses (size-abundance scaling and its temporal dynamics), we only considered every second time point to minimize temporal autocorrelation and to not artificially increase the sample size due to the interpolation.

#### S1. 4 Experimental design

**Table** S1: Overview of the experimental design

| richness | unique community compositions | replicates | experimental units | inoculum (in ml) |
|---|---|---|---|---|
| **1** | **6** | **3** | **18** | **<1.00** |
| **2** | **15** | **2** | **30** | **20.00** |
| **3** | **10** | **2** | **20** | **13.33** |
| **4** | **15** | **2** | **30** | **10.00** |
| **5** | **6** | **2** | **12** | **8.00** |
| **6** | **1** | **5** | **5** | **6.66** |

**Section S2: Predicting size-abundance scaling across temperature and richness gradients**

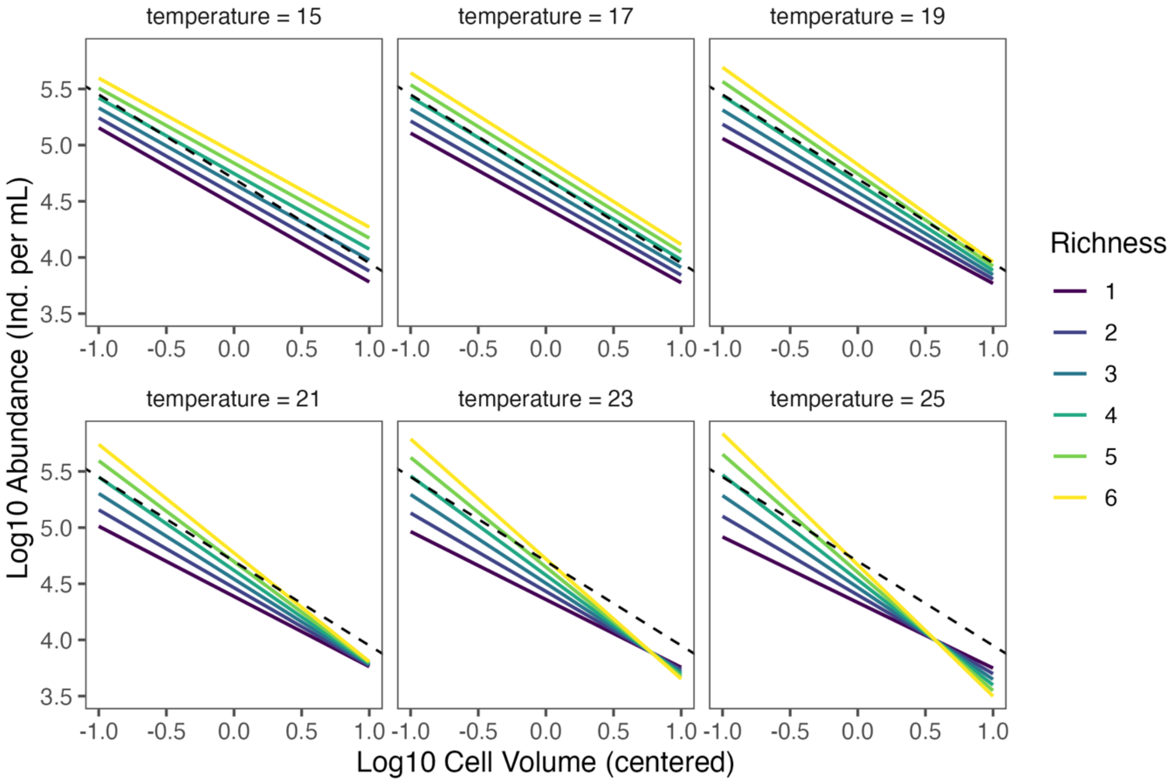


**Figure S1:** Temperature and richness interact to alter size-abundance relationships across protist communities. This figure is another visualization (see Figure 2) of the mixed model predicting total log10 abundance (number of individuals) with mean log10 cell volume (μm μL-1), temperature (15 °C, 17 °C, 19 °C, 21 °C, 23 °C, and 25 °C), and community richness (1, 2, 3, 4, 5, and 6 species), as well as interactions between those predictors based on the model presented in table 1. The dashed line indicates the expected slope of -0.75.

**Section S3: Using evenness as a proxy of realized species richness**

We experimentally manipulated initial species richness, but since community assembly and coexistence are dynamic processes where abundances change through time and extinctions may occur, we also calculated species evenness as a proxy of realized species richness. Since some species may not go extinct, but only reach very low densities, this measure is more appropriate than species richness per se. Figure S2 shows that the evenness of communities changed over the duration of the experiment. The following figure shows that there is overlap between realized richness levels that could be driven by both changes in species numbers and their relative proportions. For some communities realized richness dropped to 0 late in the experiment indicating that a single species remained and hence confirming that occasionally extinctions occurred. However, the figure also demonstrates that on average, realized richness and initial richness levels are tightly correlated and hence should provide qualitatively similar answers.

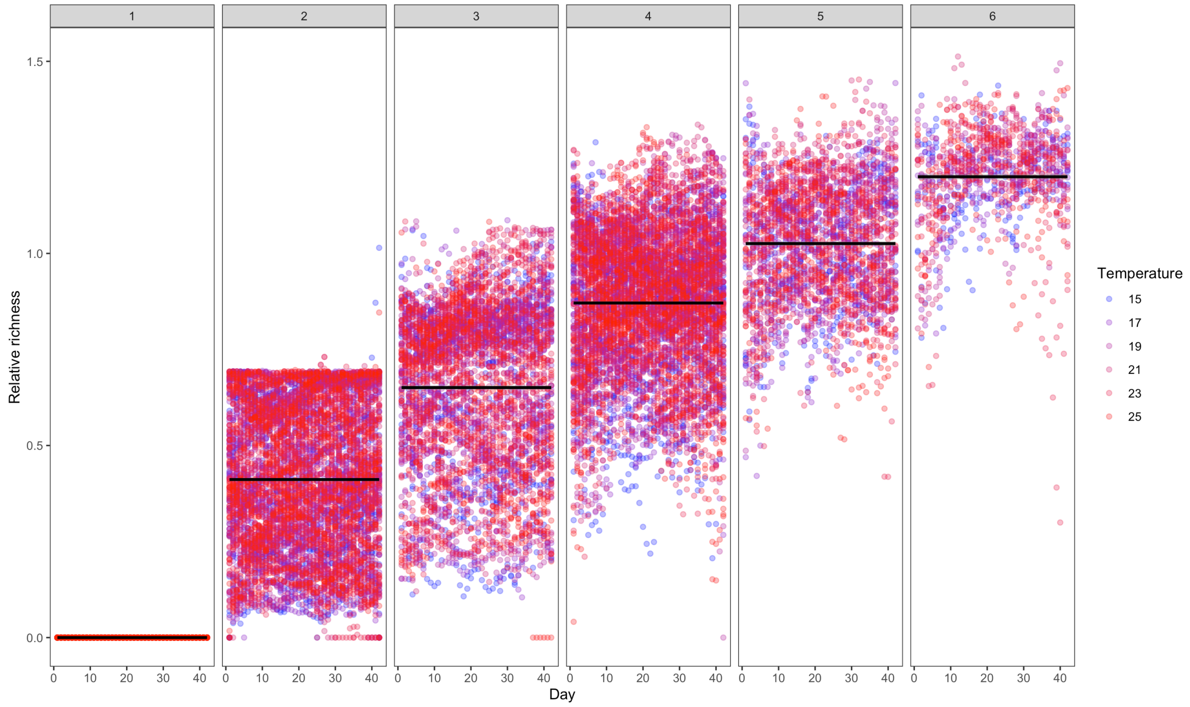


**Figure S2:** Relative richness (i.e. Shannon diversity) over the course of the experiment. The black lines show the average realized richness across the six levels of initial richness.

Subsequent analyses indeed show that using realized versus initial species richness had negligible effects on the size of the estimated richness effect and its interactions with other main effects, yielding qualitatively similar trajectories when realized species richness is used rather than initial species richness (see figure S3).

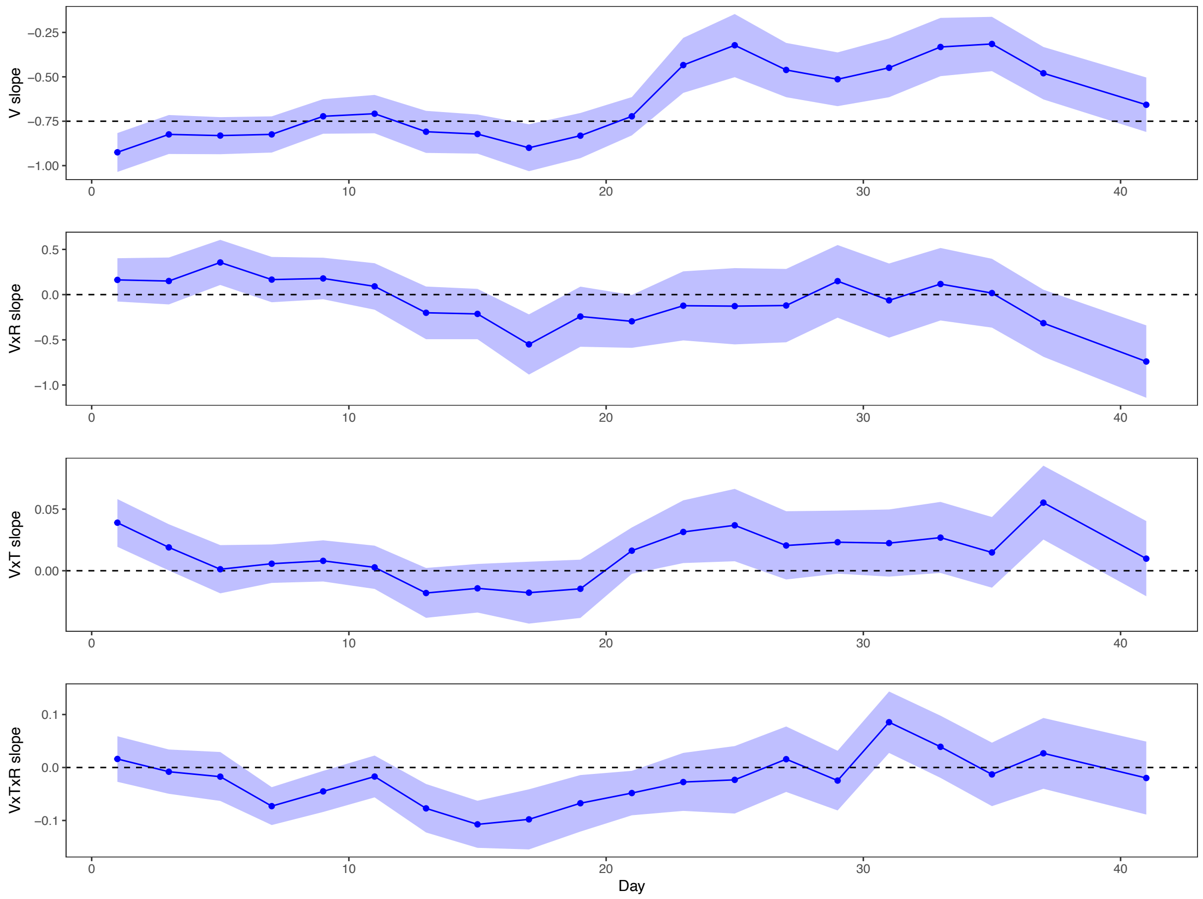


**Figure S3**: Temporal dynamics of effect sizes (and 95% confidence intervals) of a linear mixed-effects model where abundance is modelled as a function size (V = volume), temperature (= T) and realized richness (= R) and their interactions for the observed communities. The x-axis shows the day. A) Effect of cell size (V) on abundance, B-C) effect of two-way interactions between size and temperature (V x T), size and realized richness (V x R) on abundance. D) Three-way interaction between size, temperature and realized richness (V x T x R) on abundance. The dashed line indicates a slope of -0.75 in panel A and zero in panels B–D.

**Section S4: Addressing species extinction probabilities**

Since the video microscopy approach may erroneously detect species to be present at very low densities, we first define an extinction threshold of 0.01 milligram per mL for each species to assess whether species went extinct. Figure S4 shows the observed extinctions and the modelled probability to go extinct for all six species across gradients of richness and temperature. Extinction probability increases for all species at higher temperatures, but the two species going most frequently extinct in the warmest and most species rich communities are *Dexiostoma* and *Tetrahymena*. These are the two smallest species used in the experiment. This contradicts the hypothesis that steeper slopes are driven by extinctions of the largest species which have smaller population sizes and hence higher risks of going extinct (due to demographic stochasticity). In contrast, the presented data supports the conclusion that competitive asymmetries in combination with physiological responses to temperature drive the observed changes in the size-abundance slopes.

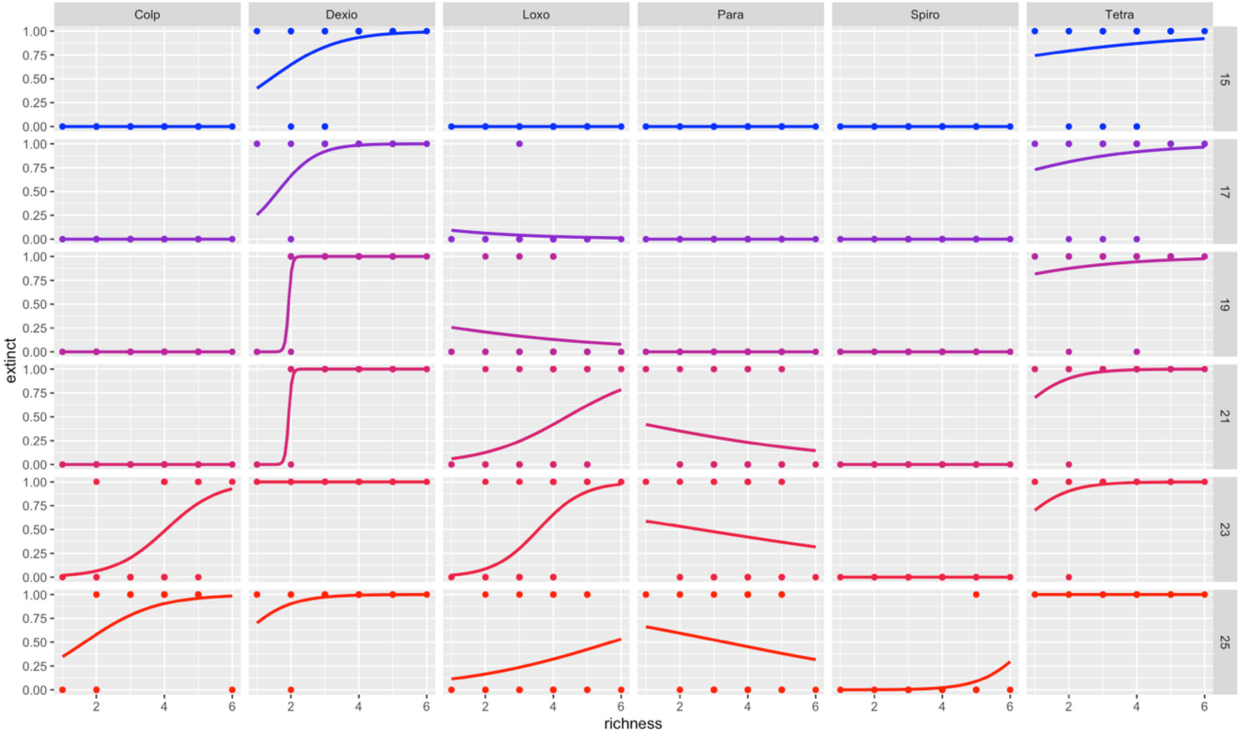


**Figure S4:** Extinction probability for the six species across temperature (columns) and richness (x-axis). Points indicate observed extinctions. Lines are the modelled probabilities of going extinct using logistic regression models for each species. The two smallest species *Dexiostoma* and *Tetrahymena* show the highest extinction probabilities across temperature and richness.

**Section S5: Interactions between size, temperature and richness through time**

Temperature and richness influenced the size-abundance relationship in distinct, time-dependent ways when comparing low- and high-richness communities. On day 11, the slope and intercept of the relationship remained stable in both low- and high-richness communities, suggesting that temperature had minimal influence on the relative abundance of small vs. large protists (i.e., no interaction, Fig. S3A). By day 17, temperature caused a steeper slope, indicating a relative increase in smaller protist abundance compared to larger ones, especially across high richness communities (i.e., negative interaction, Fig. S3C). On day 23, size-abundance scaling became shallower with higher temperatures, increasing the abundance of larger individuals compared to smaller individuals in low but not high richness communities (Fig. S3B). On day 29, this pattern was apparent in both low and high richness communities (i.e., positive interaction, Fig. S3D). These shifts highlight the dynamic influence of temperature and richness on community structure, where the balance between small and large protist abundance is regulated by time-dependent physiological and ecological processes (i.e., species interactions).

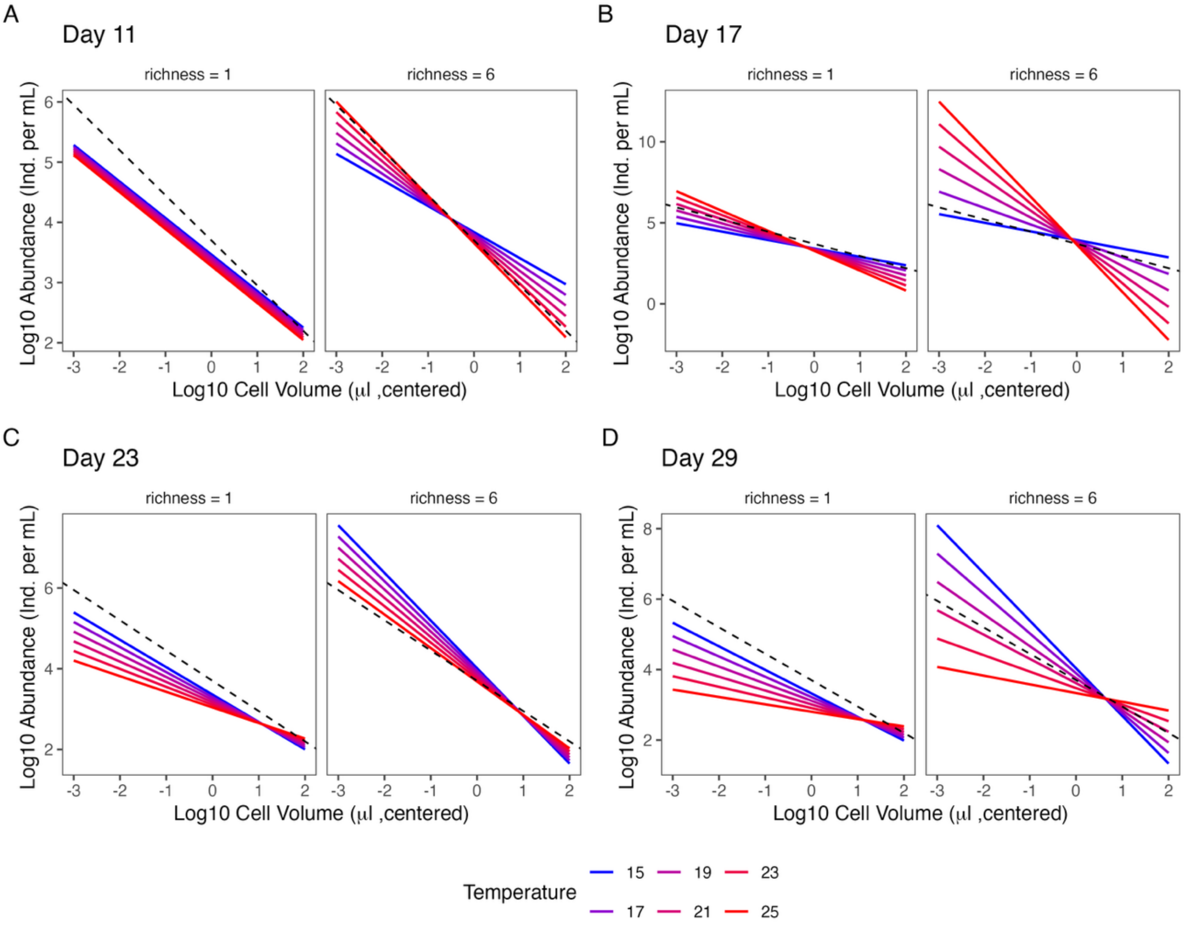


**Figure S5:** Temperature and community richness alter size-abundance relationships of protist communities across time. This figure is a visualization of the mixed model predicting total log10 abundance with mean log10 cell volume (µm $\mu L^{-1}$), temperature (15 °C, 17 °C, 19 °C, 21 °C, 23 °C, and 25 °C), and richness (low = 1 and high =

6 species number), as well as interactions between those predictors across time (A. day = 11, B. day = 17, C. day = 23 and D. day = 29; see Fig. 4D). All predictors were mean-centered for the analysis, but here the actual richness and temperature values are shown). The dashed line indicates the expected slope of -0.75.

Interpretation:

It seems that in warm environments under intense competition, smaller individuals have higher resource-use efficiency in the short-term, whereas, in the long-term, this is true for the larger ones, potentially in combination with higher survival due to the ability to store resources. Previous work has shown that intense competition selects for high resource efficiency and lower metabolism, leading larger individuals to a competitive advantage (Bierbaum *et al.* 1989; Mueller & Ayala 1981; Pianka 1970). In contrast, our study showed that this is true only in cooler conditions and the long-term. This implies that in the short-term and with increased temperature smaller individuals gain a competitive advantage due to their lower resource demands. Smaller organisms require less energy for growth, reproduction, and activity, allowing them to efficiently acquire resources and outcompete larger individuals. In contrast, larger individuals experience higher resource demands that escalate with temperature, making them less efficient. However, in the long term, larger individuals may survive better compared to smaller ones, being able to exploit available resources. This suggests that, over time, larger organisms are better able to balance resource use under elevated temperatures.
